# Dopamine Genotype Interacts with Inter-Individual Licking Received on Later-Life Licking Provisioning in Female Rat Offspring

**DOI:** 10.1101/2019.12.29.890467

**Authors:** Samantha C. Lauby, David G. Ashbrook, Hannan R. Malik, Diptendu Chatterjee, Pauline Pan, Alison S. Fleming, Patrick O. McGowan

**Author notes:** Correspondences: Alison S. Fleming, Deerfield Hall, 4th Floor, Room 4098, University of Toronto Mississauga, 3359 Mississauga Rd, Mississauga, ON L5L 1C6, Canada., Patrick O. McGowan, University of Toronto Scarborough, SW548, 1265 Military Trail, Toronto, ON M1C 1A4, Canada.

## Abstract

In most mammals, mothers exhibit natural variations in care that propagate between generations of female offspring. However, there is limited information on genetic variation that influences this propagation. We assessed early-life maternal care received by individual female rat offspring in relation to genetic polymorphisms linked to dopaminergic activity, maternal care provisioning, and dopaminergic activity in the maternal brain. We also conducted a systematic analysis of other genetic variants potentially related to maternal behavior in our Long-Evans rat population. We found that dopamine receptor 2 (rs107017253) variation interacted with the relationship between early-life maternal care received and dopamine levels in the nucleus accumbens which, in turn, were associated with later-life maternal care provisioning. We also discovered and validated new variants that were predicted by our systematic analysis. Our findings suggest that genetic variation influences the relationship between maternal care received and maternal care provisioning, similar to findings in human populations.

## 1. Introduction

The maternal environment has a substantial role in influencing offspring phenotype in adulthood, including the intergenerational transmission of maternal care through the female lineage^1^. Both maternal care received and changes in the brain that occur when the mother is preparing for postnatal offspring care involve alterations in the dopaminergic^2,3^ and oxytocinergic systems^4–6^, as well as changes in estrogen signaling^7,8^. Maternal care provisioning also involves neuropeptides aside from oxytocin, including prolactin and arginine vasopressin^9^. These mechanisms have been well-characterized in rodents and studies with humans show similar findings^10^, suggesting that the factors involved in the intergenerational transmission of maternal care are evolutionarily conserved.

Rat pups with early-life adversity induced by maternal and sibling deprivation are less attentive to foster pups and their own offspring later in life than rat pups maternally reared^11,12^. Higher licking-like tactile stimulation during the deprivation period partially mitigates the later-life impairments in maternal care provisioning^11,12^. In addition, naturally occuring variation between litters in maternal licking/grooming received can transmit between generations of offspring. High licking/grooming and low licking/grooming mothers (defined as +/-one standard deviation from the average) tend to produce female offspring that become high licking/grooming and low licking/grooming mothers to their offspring, respectively^13,14^. This effect persists in cross-fostering experiments, suggesting a prominent role of maternal behavior in the intergenerational transmission of maternal care^13^.

Previous work on the intergenerational transmission of maternal care in rats has mainly focused on proxies of maternal neglect or the tail ends of the normal distribution of maternal care received. We have more limited knowledge of factors that affect the intergenerational transmission of maternal care in the average rat mother. Seminal research examining natural variations in maternal care received between litters found that there is also substantial within-litter variation in later-life maternal care provisioning^14^. In addition, studies with human cohorts in nonclinical populations have shown a modest relationship between parental care received and parental care provisioning^15^ and found that the genotype of the individual can interact with the early-life environment or directly affect maternal care provisioning^16^. Studies in mice also show an important role of offspring genotype in maternal care received^17,18^.

Findings from our group and others have shown effects of early-life inter-individual maternal care received on later life behavior^19–23^. We have also reported gene x environment interactions involved in some of these effects^20,21^. For example, we recently reported an interaction between a dopamine receptor 2 (*Drd2*) single nucleotide polymorphism (rs107017253) and early-life average licking duration per bout (a proxy of maternal care quality) on strategy shifting and sucrose preference, two dopamine-related phenotypes^21^. Executive function measured by strategy shifting performance has previously been shown to be related to maternal care provisioning in human cohorts^24^. However, studies on the effect of genotype in the Long-Evans rat model commonly used in these experiments have been limited by the lack of knowledge of genetic variation in this outbred strain. Although there are now data available on genetic variation within wild rat populations^25^ and between inbred rat lines^26,27^, no mapping has been carried out within the Long-Evans population.

The purpose of this study was to investigate the role of genotype and gene x environment interactions in the relationship between early-life maternal licking received and later-life licking provisioning, with a specific focus on the dopaminergic system of the maternal brain. We also aimed to investigate other genes relevant to maternal behavior using a more systematic approach to identify and verify genetic variants in our Long-Evans rat population. We measured inter-individual maternal licking received within the first week of life and assessed later-life female offspring maternal licking provisioning, maternal brain dopamine (DA) and its metabolite levels, and variation in single nucleotide polymorphisms (SNPs) in genes involved in DA, estrogen, and neuropeptide function. We predicted that higher early-life licking received would correspond to higher later-life licking provisioning in female offspring. In addition, we hypothesized that this relationship would be indirectly affected by dopaminergic activity in the maternal brain and interact with female offspring genotype. The hypothesized moderated-mediation model is displayed in Figure 1.

**Figure 1.**
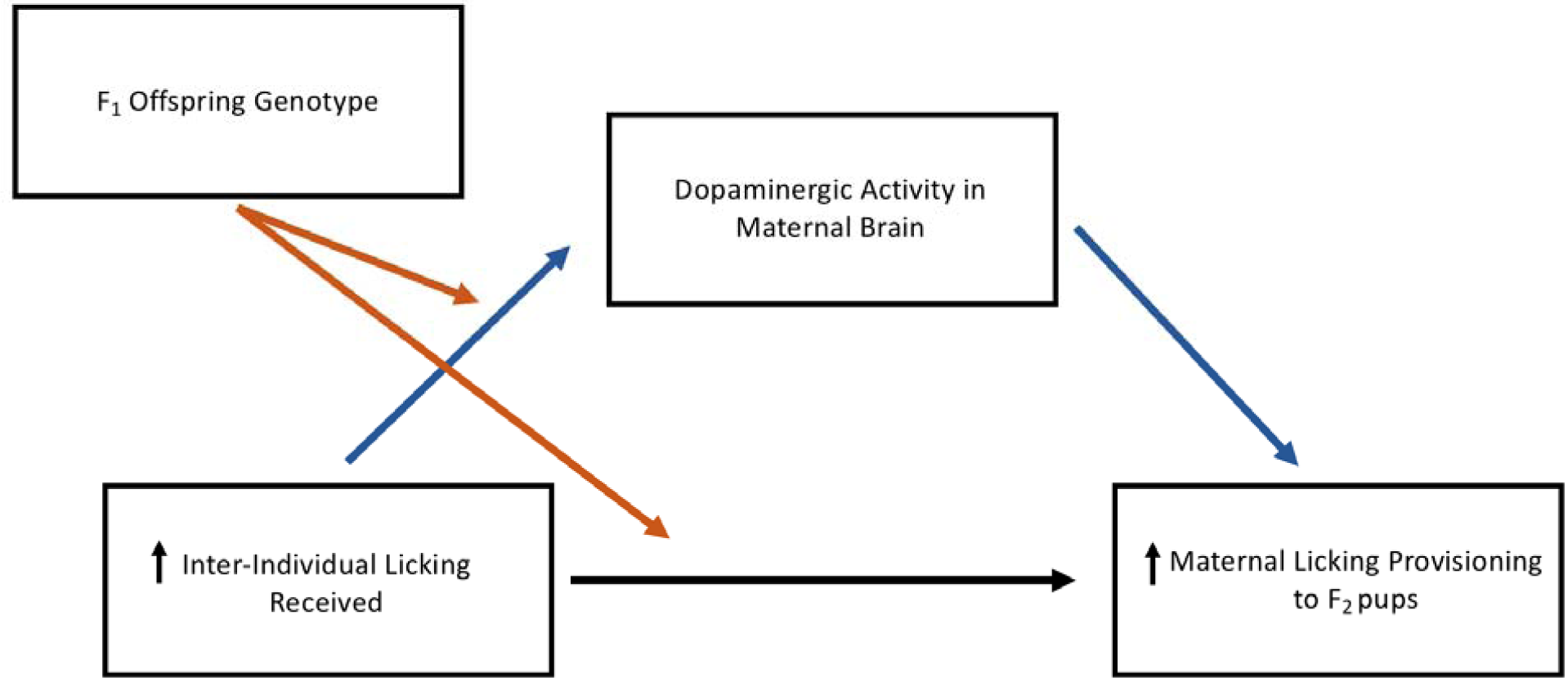
Hypothesized moderated mediation between early-life licking received and later-life licking provisioning. Inter-individual licking received early in life would positively associate with later-life maternal licking provisioning. Differences in dopaminergic activity in the maternal brain would account for this association and mediate the relationship between licking received and licking provisioning indirectly. Finally, offspring genotype in dopamine-related genes would interact with early-life inter-individual licking received and moderate later-life maternal licking provisioning and dopaminergic activity in the maternal brain.

## 2. Material and Methods

### 2.1 Rat Breeding

Seven-week-old female (n = 24) and male (n = 6) Long-Evans rats were obtained from Charles River Laboratories. They were housed in same-sex pairs on a 12:12 hour light-dark cycle (lights on at 7:00) with ad libitum access to standard chow diet and water. For breeding, one male was housed with two females for one week. Females were then housed separately and weighed weekly throughout pregnancy. The breeding males were used multiple times to produce four cohorts of litters in this study. All animal procedures were approved by the Local Animal Care Committee at the University of Toronto Scarborough and conformed to the guidelines of the Canadian Council on Animal Care.

The pregnant F_0_ females were checked for parturition starting three weeks after breeding at 9:00 and 17:00. Postnatal day (PND) 0 was determined if the birth occurred between 9:00 and 17:00 or if pups were noted at 9:00 but have not nursed yet. Pups found at 9:00 with a visible milk band were determined to be PND 1. At PND 1, litters were culled to five to six female pups and individually weighed. We focused on smaller litters in order to accurately measure maternal care received. Therefore, male offspring were not examined for this study. A total of 136 F_1_ female pups were assessed for maternal care received.

### 2.2 Maternal Care Received Observations

At PND 1, 3, 5 and 7, maternal care was assessed as previously reported^21^. From 10:00 to 17:00 in the light phase, litters were briefly separated from their mother and individually marked using odorless and tasteless food colouring (Club House, London, Canada) to distinguish between siblings. The entire litter was then placed in the opposite corner of the established nest and maternal behavior was observed for 30 minutes using Observer XT 11.5 (Noldus Information Technology, Wageningen, The Netherlands). To establish inter-rater reliabilities on behavioral observations, three researchers coded the same mothers with an experienced coder until high reliability (>90%) was consistently met.

The order pups were retrieved was recorded manually. Duration and frequency of anogenital licking and body licking were coded for individual pups, meaning each pup had designated keys in the Observer software. Hovering, nursing (blanket and arched-back), and nest-building were coded as a litter, since these behaviors typically involve whole litters. Self-directed behaviors (feeding and self-grooming) by the mother were also coded. Cages were not changed throughout the maternal behavior observation period.

Total duration of licking (anogenital and body licking) and average duration of a lick bout (total duration of licking/number of licking bouts) across all four observation days for each pup were calculated as measures of maternal care. This was referred as “total licking duration” and “average licking duration”, respectively.

Female offspring were weaned at PND 22 and were pair-housed with siblings. They were weighed periodically until adulthood (PND 75).

### 2.3 Maternal Care Provisioning Observations

A subset of adult F_1_ female offspring (n = 54) were bred with sexually experienced males (n = 6) for one week. Six females were unable to get pregnant after two rounds of breeding and two females failed to lactate following parturition, with a total of 46 F_1_ rat mothers being used for intergenerational maternal care analysis. Births were checked starting three weeks after breeding at 9:00 and 17:00. At PND 1, cages were changed with no culling of pups. The litter sizes ranged from 4 to 18 pups and were not significantly correlated with maternal licking provisioning (Pearson’s r = 0.001, p = 0.994).

From postnatal days 2 to 9, each litter was video recorded for one hour three times during the light phase (9:00-10:00, 13:00-14:00, 17:00-18:00) and three times during the dark phase (21:00-22:22, 1:00-2:00, 5:00-6:00) using security cameras connected to a DVR system (Swann Communications Ltd.). These videos were coded with Observer XT 10.5 (Noldus Information Technology) for maternal behavior by four coders who were blind to the genotype of the F_1_ rat mother. Nursing, licking, nest-building, and other self-directed behaviors were scored every three minutes based on previous literature^14^. A total of 120 observations per F_1_ mother per day were coded and maternal licking provisioning was represented as a percentage of the frequency of licking behavior (body and anogenital combined) coded over total observations.

After 10:00 on PND 9 of the F_2_ pups, the F_1_ rat mother was separated from her pups and was sacrificed with CO_2_ inhalation and decapitation. Liver and whole brain tissue were collected and placed in dry ice or flash-frozen in isopentane, respectively. Tissue was stored in −80°C until further processing.

### 2.4 High Performance Liquid Chromatography

Forty-six maternal rat brains were sliced and microdissected using a Leica CM3050S cryostat (Leica Microsystems, Wetzlar, Germany). Medial prefrontal cortex (+4.20 mm to +2.70 mm Bregma), nucleus accumbens core and shell (+2.20 mm to +1.20 mm Bregma), medial preoptic area (0.30mm to −0.80mm Bregma), dorsal hippocampus (control brain region; −2.30 mm to - 3.30 mm Bregma), and the ventral tegmental area (−5.20 mm to −5.60 mm Bregma) were identified using an adult rat brain atlas^28^. 5 μL of 1.0 M ascorbic acid was added to each sample to stabilize the neurotransmitters followed by storage at −80°C.

To prepare the samples, the brain tissue was thawed on ice, suspended in 20 μl artificial cerebrospinal fluid (Harvard Apparatus, Holliston, MA) and homogenized by four pulses of sonication (2 seconds per pulse). 2 μl of brain homogenate from each sample was analyzed for protein concentration using BioRad protein assay reagent (BioRad, Hercules, CA). 1 μl of 0.2 M perchloric acid per sample was added to the remaining homogenate and was centrifuged at 10,000 rpm for 10 minutes at 4°C. The supernatant was collected and stored at −80°C.

Dopamine (DA) and 3, 4-dihydroxyphenylacetic acid (DOPAC) levels were measured by high performance liquid chromatography (HPLC) as described by previous studies^21,29^. A BAS 460 MICROBORE-HPLC system with electrochemical detection (Bio-analytical Systems Inc., West Lafayette, IN) was used together with a Uniget C-18 reverse phase microbore column (#8949, Bio-analytical Systems Inc.). The mobile phase contained a buffer [0.1 M monochloro acetic acid, 0.5 mM Na-EDTA, 0.15 g/L Na-octylsulfonate and 10 nM sodium chloride, pH 3.1], acetonitrile and tetrahydrofuran at a ratio of 94:3.5:0.7. The flow rate was 1.0 ml/min, and the working electrode (Uniget 3 mm glassy carbon, BAS P/N MF-1003) was set at 550 mV vs. Ag/Ag/Cl reference electrode. Detection gain was 1.0 nA, filter was 0.2 Hz, and detection limit was set at 20 nA. 5 μl of the sample supernatant was directly injected into the column. External standards for DA and DOPAC (Sigma Aldrich, St. Louis, MO) of known concentrations were used to quantify and identify peaks on the chromatogram. Under these parameters, the retention times for DA and DOPAC were approximately 3.7 minutes and 5.5 minutes, respectively.

DA and 3, 4-dihydroxyphenylacetic acid (DOPAC) levels were normalized against total protein concentration for each sample. DOPAC divided by DA, or the DOPAC/DA ratio, was calculated as a measure of dopaminergic activity in each brain area. We were unable to analyze two prefrontal cortex samples, two nucleus accumbens samples, one medial preoptic area sample and three ventral tegmental area samples due to lost tissue during collection.

### 2.5 Genotyping

To identify variants which segregate within the Long-Evans population, we took advantage of large amounts of RNA-seq data available online in the population. We identified 119 RNA-seq datasets on the Gene Expression Omnibus (GEO; Supplementary Table 1) which we could use for variant calling. Fastq files were downloaded, aligned and variants called following the Genome Analysis Toolkit (GATK) guidelines. In brief, adaptors were trimmed from the fastq files using trim galore (version 0.4.1). The aligned files were aligned to the rn6 rat reference genome using STAR (version 2.6.0.), using two-pass mode^30^. Read groups were added and duplicates were marked using picard tools. NCigar reads were split using GATK (version 4.0.8.1), and variants were called using HaplotypeCaller. Joint-calling was carried out using all 119 samples using GenomicsDBImport and GenotypeGVCFs. Variant recalibration was carried out, using a ‘true positive’ variant list from the rat genome database. This provided us with a list of high quality single nucleotide polymorphisms (SNPs) and small INDELs.

**Table 1.**
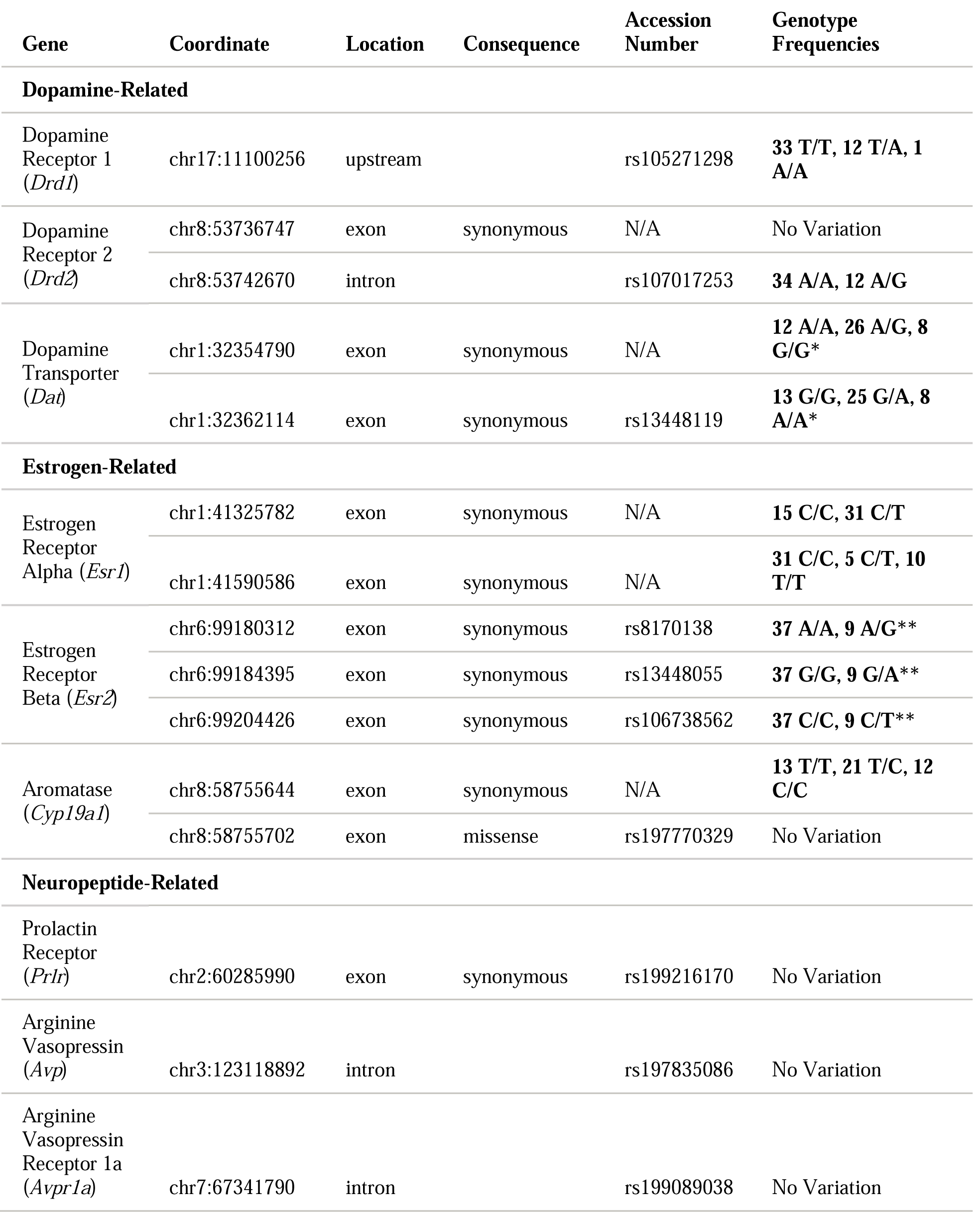
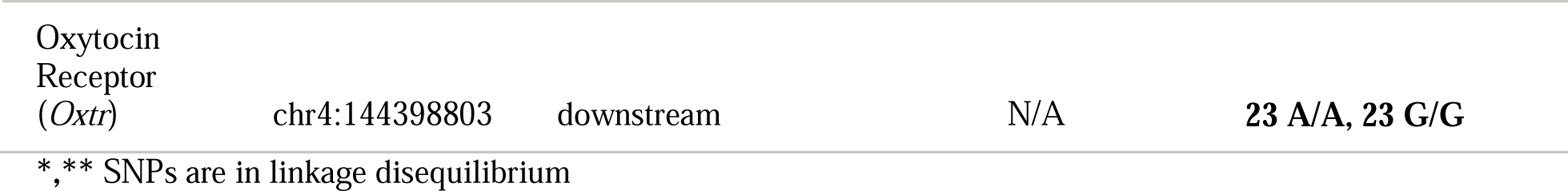
List of candidate single nucleotide polymorphisms assessed for variation, information related to their location and function, and genotype frequencies in our Long-Evans rat population.

To validate variation in candidate SNPs, liver DNA from the F_1_ rat mothers was extracted using an EZNA Tissue DNA extraction kit (Omega Bio-tek, Norcross, GA) and assessed for SNPs in genes relevant to maternal behavior, broadly including DA-related, estrogen-related, and neuropeptide-related genes (Table 1). Purified DNA (35 ng/μl) was submitted for a multiplex assay at The Centre for Applied Genomics (SickKids, Toronto, Canada).

### 2.6 Statistical Analysis

All statistical analyses were performed using SPSS (IBM Corporation). To examine the relationship between early-life maternal licking received and later-life maternal licking provisioning, a Pearson correlation was used between total or average licking duration received and the percentage of licking provisioning observed. In addition, to examine the relationship between later-life maternal licking provisioning and dopaminergic activity, a Pearson correlation was used between percentage of licking provisioning observed and the DOPAC/DA ratio in each brain area, with a false discovery rate (FDR) correction using the Benjamini-Hochberg procedure to account for multiple analyses. Significant correlations with DOPAC/DA ratio in a brain area were followed up with Pearson’s correlations with DOPAC and DA levels separately. To examine the effects of genotype, a one-way ANOVA was used to compare offspring with each varying genotype to early-life maternal licking received and later-life maternal licking provisioning with a FDR correction to account for multiple SNP analyses for each outcome licking measure. Significant effects of genotype were followed with a Tukey’s post-hoc test. To examine gene x environment interactions and the mediating role of dopaminergic activity on later-life maternal licking provisioning, we used Hayes PROCESS module for SPSS (Version 3.2) using a simple moderation (Model 1) with a FDR correction to account for multiple analyses for each outcome dependent variable and a moderated mediation (Model 7)^31^. A moderation tests for an interaction between an independent variable (*X*) and a moderator variable (*W*) on a dependent variable (*Y*) and a mediation tests for indirect associations of the independent (*X*) and dependent variable (*Y*) by a third causal variable (the mediator *M*_*i*_). PROCESS is a flexible modelling module that can conduct moderation and mediation analyses using multiple regression and conduct post-hoc analyses of the conditional effects of a focal moderator and the indirect effects of a mediator.

The statistical model used for a simple moderation (Model 1) was:

Conditional direct effects of *X* on *Y* = *b*_*1*_ + *b*_*3*_*W*

The statistical model used for a moderated mediation (Model 7) was:

Conditional indirect effects of *X* on *Y* thorough *M*_*i*_ = (*a*_*1i*_ + *a*_*3i*_*W*)*b*_*i*_

Direct effect of *X* on *Y* = *c’*

Multi-categorical moderator variables (i.e., more than two genotypes) were analyzed using the indicator coding system. All effects were considered statistically significant if p ≤ 0.05 and marginally significant if p ≤ 0.10.

## 3. Results

### 3.1 Relationship between Early-Life Licking Received and Later-Life Licking Provisioning

Total licking duration received across all maternal care observation periods (PND 1, 3, 5, and 7) averaged 168.79 ± 18.6 seconds and ranged from 0 to 450.72 seconds (Figure 2A). Average licking duration per bout of licking received averaged 8.02 ± 0.56 seconds and ranged from 0 to 13.93 seconds (Figure 2B).

**Figure 2.**
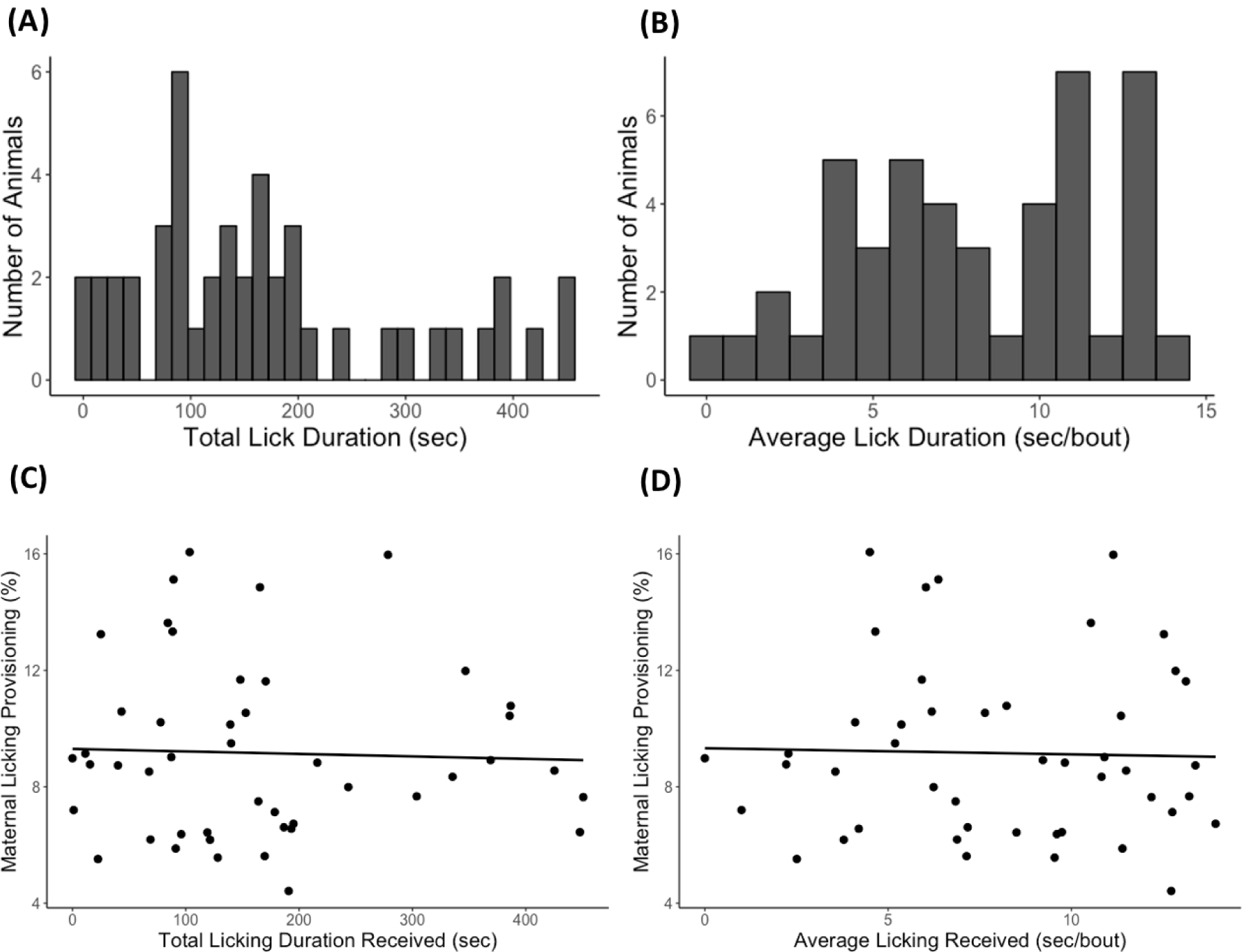
The distribution of maternal licking received within the first week of life and its direct association with later-life maternal licking provisioning. (A) Total licking duration (15-second bins) and (B) average licking duration per bout (1-second bins) received across all observation days (PND 1, 3, 5, and 7) for the female offspring tested (n = 46). There were no significant correlations between (C) total licking duration and (D) average licking duration per bout with later-life licking provisioning.

We first examined the relationship between early-life maternal care received and later-life maternal care provisioning. There were no direct associations between early-life licking received and later-life licking provisioning. Neither total licking received (Pearson’s r = −0.036, p = 0.812; Figure 2C) nor average licking received (Pearson’s r = −0.027, p = 0.861; Figure 2D) significantly correlated with later-life licking provisioning.

### 3.2 Relationship between Early-Life Licking Received, Later-Life Licking Provisioning and Dopaminergic Activity in the Maternal Brain

We then examined relationships between maternal care received and provisioning with DOPAC/DA ratio in the medial prefrontal cortex, nucleus accumbens, medial preoptic area, and ventral tegmental area in the F_1_ maternal brain. There was a significant negative correlation between later-life licking provisioning and DOPAC/DA ratio in the nucleus accumbens (Pearson’s r = −0.389, FDR adjusted p = 0.036; Figure 3A). Upon further analysis of DOPAC and DA levels separately, there was a significant positive correlation between later-life licking provisioning and DA levels (Pearson’s r = 0.346, p = 0.021; Figure 3B) and no correlation with DOPAC levels (Pearson’s r = −0.060, p = 0.701; Figure 3C) in the nucleus accumbens. There were no other associations between later-life licking provisioning and DOPAC/DA ratio in any other brain regions.

**Figure 3.**
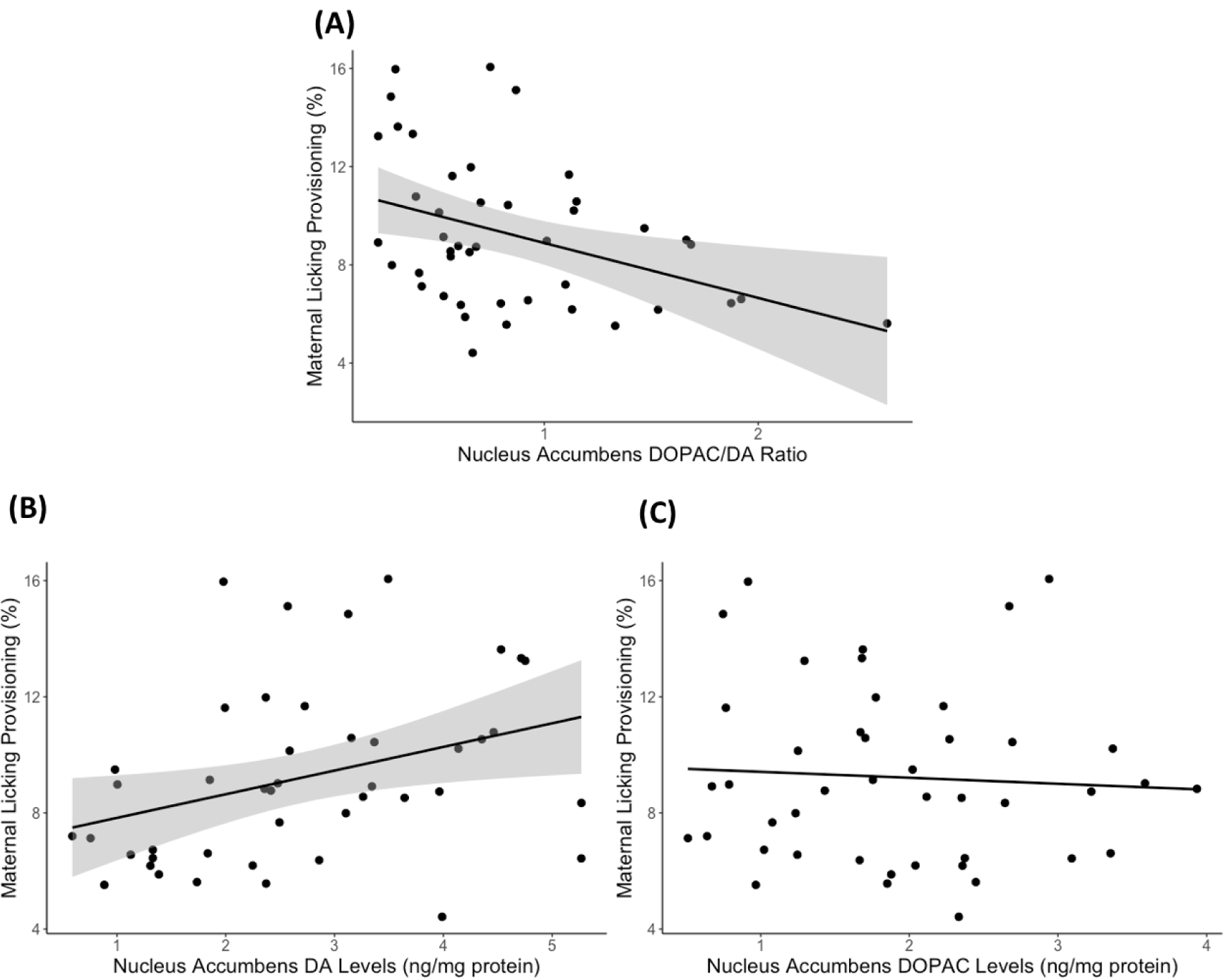
Dopaminergic activity in the nucleus accumbens of the maternal brain is associated with later-life licking provisioning. (A) The DOPAC/DA ratio in the nucleus accumbens was negatively correlated with later-life licking provisioning from postnatal day 2-9. Upon further analyses, the (B) dopamine (DA) levels in the nucleus accumbens was positively correlated with later-life licking provisioning and (C) there was no correlation with DOPAC levels in the nucleus accumbens. Scatterplots are displayed with 95% confidence interval in grey for the DOPAC/DA ratio and DA levels in the nucleus accumbens.

### 3.3 Effects of Genotype

We detected 792,253 SNPs which segregate in the Long-Evans population. These variants are available as a UCSC genome browser track at http://genome.ucsc.edu/cgi-bin/hgTrackUi?hgsid=832959263_fmyKPx8JTUe0A2koSyaIFEUT1jGA&c=chr1&g=hub_1969047_Long_Evans_vcf. We chose seven genes relevent to maternal care provisioning, and examined 16 variants within these genes.

Of the 16 variants in genes relevant to maternal care provisioning detected, 11 showed variation in our animals (Table 1). These SNPs were in dopamine receptor 1 (*Drd1*; rs105271298), dopamine receptor 2 (*Drd2*; rs107017253), dopamine transporter (*Dat*; chr1:32354790 and rs13448119), estrogen receptor alpha (*Esr1*; chr1:41325782 and chr1:41590586), estrogen receptor beta (*Esr2*; rs8170138, rs13448055, and rs106738562), aromatase (*Cyp19a1*; chr8:58755644), and oxytocin receptor (*Oxtr*; chr4:144398803). We did not detect variation in the SNPs we tested in prolactin receptor (*Prlr*), arginine vasopressin (*Avp*) and arginine vasopressin receptor 1a (*Avpr1a*).

The one female offspring with the *Drd1* A/A genotype was combined with the A/T genotype for analysis. In addition, since the three SNPs within *Esr2* and the two SNPs within *Dat* were in complete linkage, only one SNP in each gene (rs8170138 and rs13448119, respectively) was used for analysis.

There was a main effect of *Oxtr* (chr4:144398803) genotype (F _(1,44)_ = 9.14, FDR adjusted p = 0.032) and a marginal effect of *Esr2* (rs8170138) genotype (F _(1,44)_ = 6.78, FDR adjusted p = 0.056) and *Cyp19a1* (chr8:58755644) genotype (F _(2,43)_ = 4.23, FDR adjusted p = 0.056) on average licking received. For *Oxtr*, homozygous G/G female offspring received lower average licking per bout than homozygous A/A female rat offspring (Figure 4A). In addition, there was a main effect of *Esr1* (chr1:41590586) genotype (F _(2,43)_ = 9.54, FDR adjusted p = 0.002) on later-life licking provisioning. Heterozygous C/T female rat offspring provided more licking than homozygous C/C (Tukey’s post-hoc p = 0.001) and T/T (Tukey’s post-hoc p = 0.001) female offspring (Figure 4B). There were no main effects of DA-related SNPs (*Drd1, Drd2* or *Dat*) on early-life maternal licking received or provisioning.

**Figure 4.**
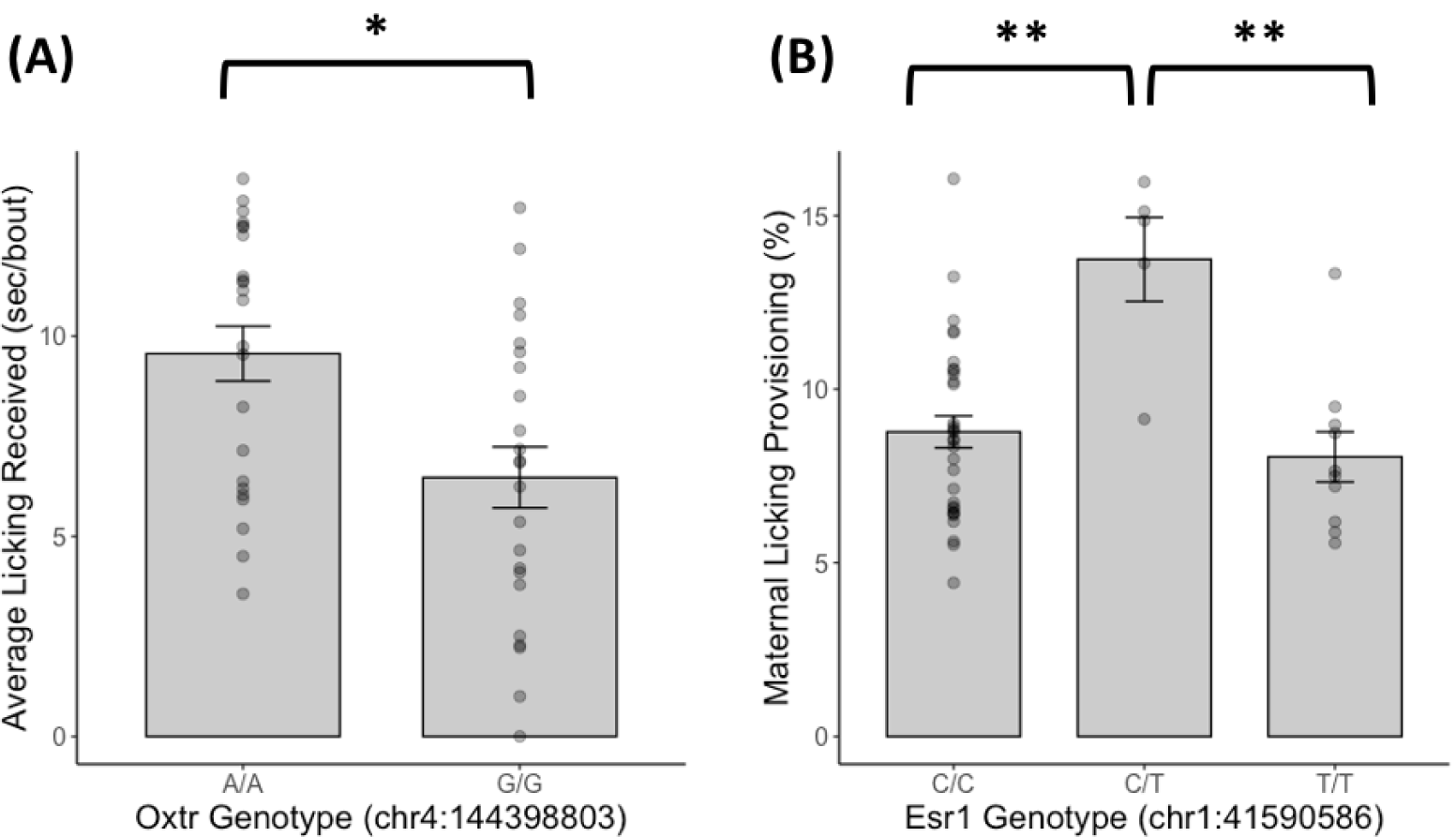
Offspring genotype within the oxytocin receptor and estrogen receptor alpha genes affect early-life licking received or later-life licking provisioning respectively. (A) Homozygous G/G female rat offspring for oxytocin receptor (*Oxtr*; chr4:144398803) received lower average licking per bout than homozygous A/A female rat offspring and (B) heterozygous C/T female rat offspring for estrogen receptor alpha (*Esr1*; chr1:41590586) provided more licking than homozygous C/C and T/T female rat offspring. Barplots are displayed with mean +/-standard error with individual data points. * p < 0.05, ** p = 0.001 with Tukey’s post-hoc tests.

### 3.4 Gene x Environment Interactions

We restricted our gene x environment analyses to dopamine-related SNPs (*Drd1, Drd2, Dat*) because we hypothesized that the indirect relationship between early-life licking received and later-life licking provisioning would be mediated by dopaminergic activity in the maternal brain. We also restricted our outcome measures to later-life licking provisioning and DA levels in the nucleus accumbens because this was the only dependent variable that significantly correlated with later-life maternal licking provisioning. We analyzed both total licking received and average licking received as independent variables.

*Drd2* (rs107017253) genotype interacted with average licking received on DA levels in the nucleus accumbens (Coefficient β = −0.2643 ± 0.093, t _(1,42)_ = −2.8366, FDR adjusted p = 0.0426; Figure 5). More specifically, there was a significant effect for the A/G genotype (Effect = 0.230 ± 0.071, t = 3.2363, p = 0.0024) but not the A/A genotype (Effect = −0.0344 ± 0.060, t = −0.5700, p = 0.5718). No other gene x environment interactions were observed with total licking received or other dopamine-related SNPs. In addition, there were no significant gene x environment interactions on later-life licking provisioning.

**Figure 5.**
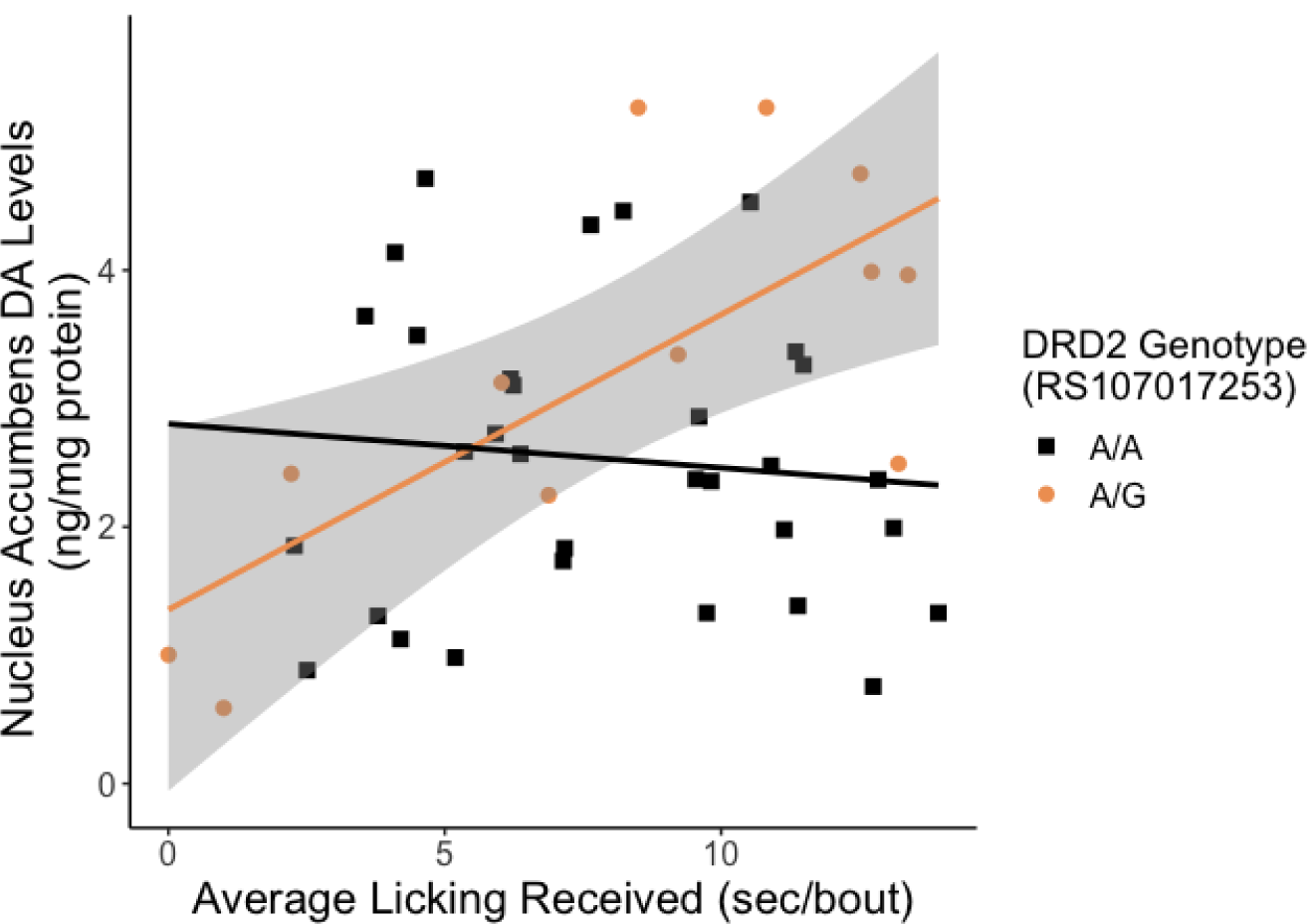
The association between average licking received and nucleus accumbens dopamine (DA) levels in the maternal brain is moderated by dopamine receptor 2 (*Drd2*) genotype. Heterozygous A/G female rat offspring had higher levels of DA in the nucleus accumbens with higher early-life average licking received. Homozygous A/A offspring do not show an association with early-life average licking received. Scatterplots are displayed with regression lines for A/A (black) and A/G (orange) with the 95% confidence interval for the A/G genotype in grey.

### 3.5 Moderated Mediation of Early-Life Licking Received on Later-Life Licking Provisioning

Based on the reported results, we conducted a moderated mediation analysis in PROCESS (Model 7) using average licking received as the independent variable, percent licking provisioning as the outcome dependent variable, DA levels in the nucleus accumbens as a mediator, and *Drd2* (rs107017253) genotype as a moderator. Figure 6 displays the moderated mediation statistical model with all output coefficient β values. These β values also reflected previous analyses with a statistically significant *Drd2* (rs107017253) genotype by average licking received interaction on DA levels in the nucleus accumbens as well as a statistically significant positive correlation between DA levels in the nucleus accumbens and maternal care provisioning.

**Figure 6.**
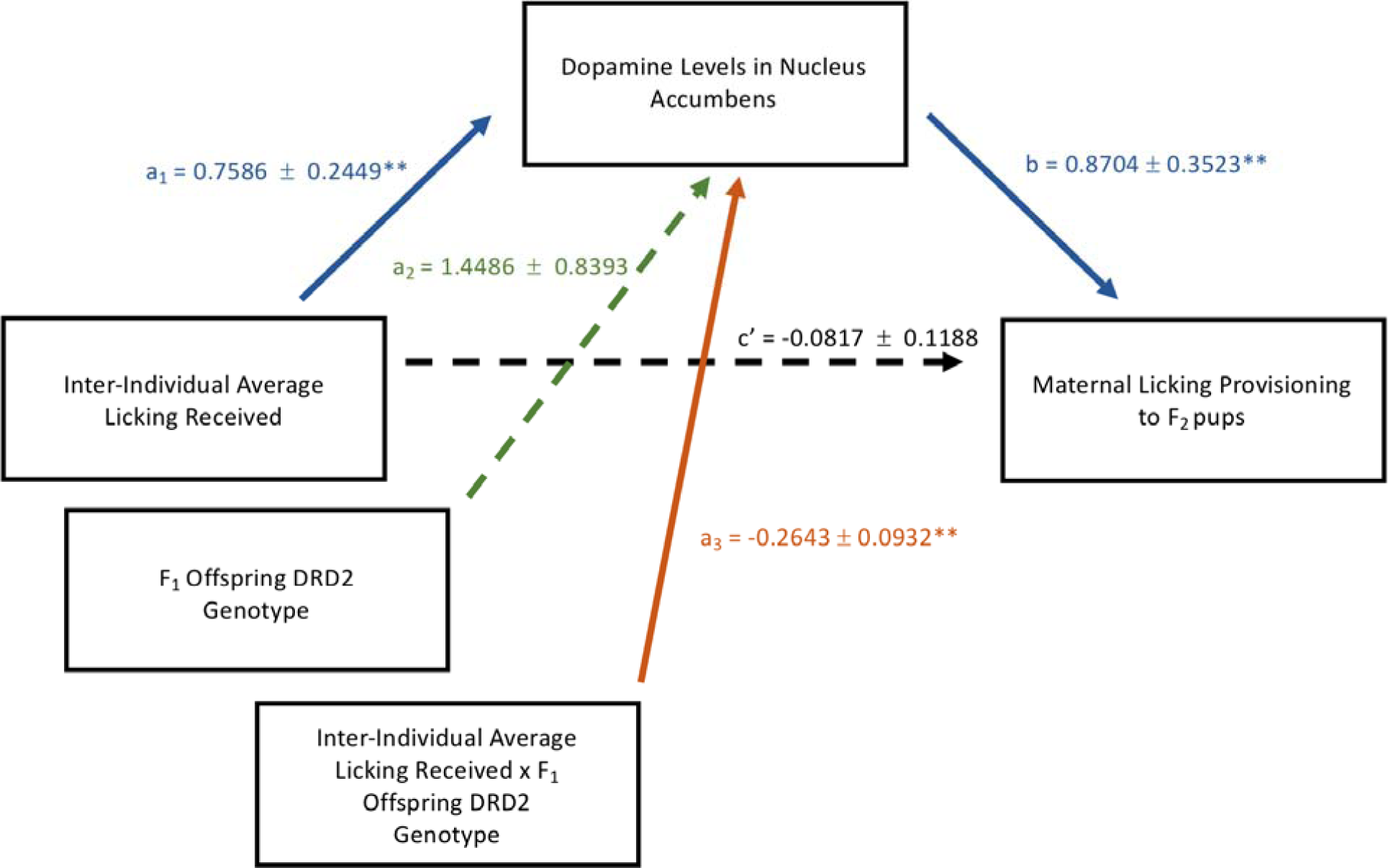
The moderated mediation statistical model (Model 7) analyzed with PROCESS with all output coefficient β values. There was a significant moderation of *Drd2* (rs107017253) and average licking received (solid orange) on dopamine levels in the nucleus accumbens. Dopamine levels in the nucleus accumbens, in turn, significantly mediated (solid blue) the relationship between average licking received and maternal licking provisioning to the F_2_ pups. ** p < 0.05

We found an indirect moderation of *Drd2* genotype between average licking received and later-life licking provisioning by DA levels in the nucleus accumbens (Index of moderated mediation = −0.2301, Bootstrap 95% CI [-0.4697, −0.0193]). Specifically, there was a significant indirect effect for the A/G genotype (Effect = 0.2002, Bootstrap 95% CI [0.0185, 0.4063]) but not the A/A genotype (Effect = −0.0299 Bootstrap 95% CI [-0.1533, 0.0910]). This suggests that higher early-life average licking received was associated with higher DA levels in the nucleus accumbens of the maternal brain but only in female rat offspring with the A/G *Drd2* genotype. Higher DA levels in the nucleus accumbens of the maternal brain of the female rat offspring with the A/G *Drd2* genotype was then associated with higher later-life licking provisioning.

## 3. Discussion

In this study, we investigated the role of genotype in the transmission of inter-individual maternal care across generations of female rat offspring. To our knowledge, this is the first study that has examined the relationship between inter-individual maternal care received and maternal care provisioning to the next generation of offspring and the genetic factors that could influence this relationship. We found that the relationship between early-life licking received and later-life licking provisioning was indirectly affected by DA levels in the nucleus accumbens and dependent on *Drd2* genotype. More specifically, female rat offspring with the A/G genotype showed a positive relationship between average licking received and dopamine levels in the nucleus accumbens of the maternal brain; there was no relationship with female rat offspring with the A/A genotype. The higher DA levels in the nucleus accumbens corresponded with higher maternal licking provisioning from postnatal day 2-9. The updated moderated-mediation model is displayed in Figure 7. In addition, estrogen-and oxytocin-related SNPs affected both early-life licking received and later-life licking provisioning, suggesting that genotype can interact with the early-life maternal environment and influence later-life maternal care provisioning. Given that the early-life licking received we observed represented a continuous range of maternal care, these results suggest that genotype has an important role in the transmission of maternal care across generations in the average rat mother.

**Figure 7.**
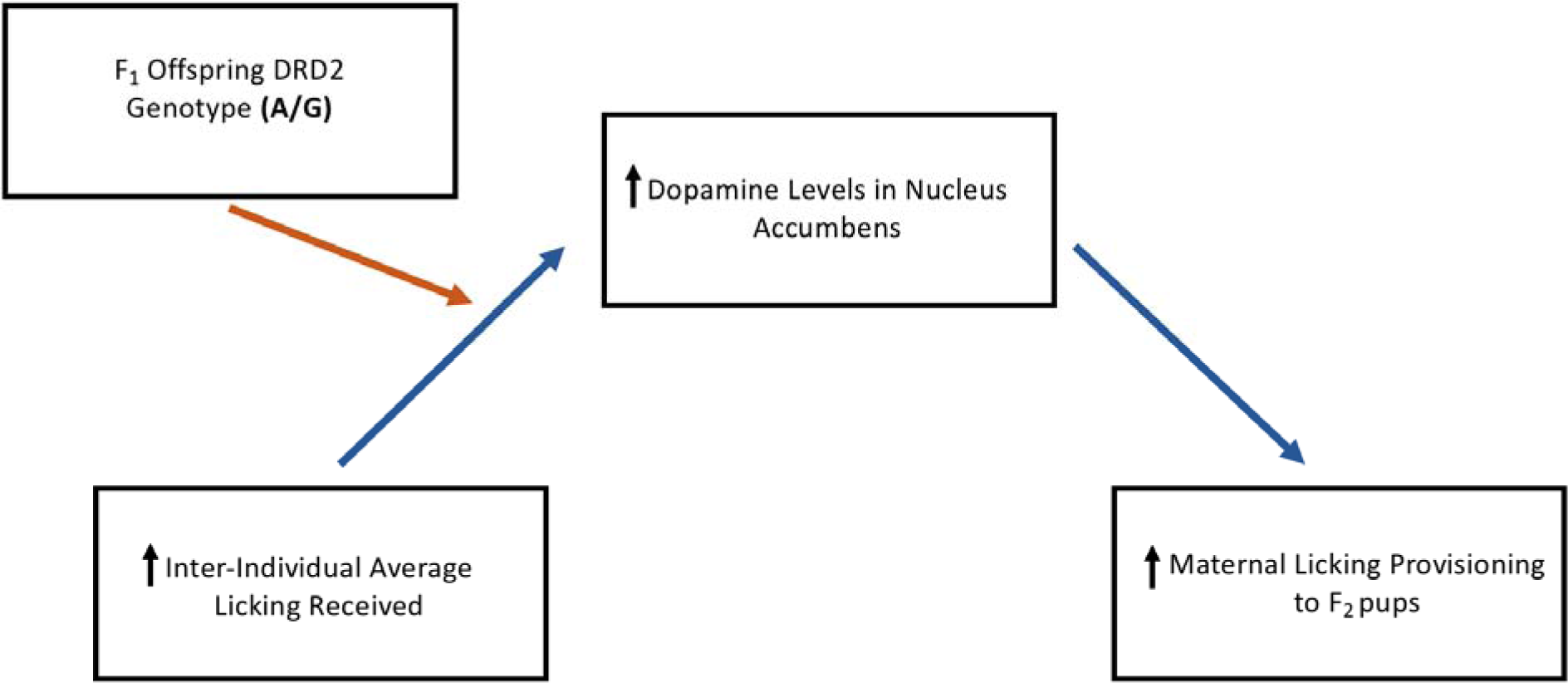
Updated moderated mediation model between early-life licking received and later-life licking provisioning. Offspring *Drd2* genotype moderated the indirect relationship between early-life average licking received and later-life maternal licking provisioning by dopamine levels in the nucleus accumbens.

We first investigated the direct relationship between early-life licking received and later-life licking provisioning and did not find any correlation between the two measures. Previous work in rats has shown that inducing early-life adversity and, in studies observing natural variations in maternal care, the outliers of maternal care received (+/-one standard deviation) can transmit across generations of offspring^11,13,14^, demonstrating that early-life environmental influences are sufficient to alter maternal care to the next generation. However, there is substantial within-litter variation in the intergenerational transmission of maternal care, especially with female offspring born to mothers that provide average levels of licking/grooming^14^. Since we did not select for the tail ends of the normal distribution in inter-individual licking received across all litters for the maternal care provisioning observations, our findings may reflect the substantial variation in intergenerational maternal care transmission within the normal distribution of maternal care received.

We found that the DOPAC/DA ratio (a measure of dopaminergic activity) in the nucleus accumbens was negatively associated with later-life licking provisioning, mainly due to the positive correlation with DA levels than a negative correlation with DOPAC levels. Previous studies have shown links between maternal care provisioning and dopaminergic activity in the nucleus accumbens, especially the nucleus accumbens shell^3,32–34^. The relationship between early-life licking received and later-life licking provisioning was also mediated by levels of dopamine in the nucleus accumbens in the maternal brain, suggesting that the nucleus accumbens may be an important site of intergenerational transmission of maternal care. Artificially reared female rat pups show later-life elevated basal DA levels and lower pup-evoked DA levels in the nucleus accumbens and a reduction in maternal care^11,34^ than normal maternally reared offspring. Additional studies are needed to explore the potential role of the nucleus accumbens and its subregions in the transmission of the effects of maternal care received at the early-life pup stage.

We detected 792,253 genetic variants that appear to segregate in the Long-Evans population, overlapping 20,508 Ensembl gene annotations. This number is likely an underestimation due to RNA-seq data being enriched in reads from transcribed regions. Of sixteen variants of interest that we verified using massarray, five were not polymorphic. The two most likely reasons for this are that either a) the variant didn’t exist in our population of Long-Evans, either due to genetic drift, different suppliers, or simple change; or b) they were false positive calls, due to the greater difficulty in calling variants from RNA-seq data than from DNA-seq data. However, the majority of our calls variants were confirmed (69%), and five are novel variants (83% of six novel variants probed). These findings show that this method is useful in understudied populations, which is important because a large amount of rat genetic variation is still undocumented. The data we have generated and publically shared on the genetic variation in the Long-Evans rat strain stands to be a useful resource for other researchers studying the effects of natural genetic variation and epigenetic mechanisms on phenotype.

We found that single nucleotide polymorphisms (SNPs) in candidate genes related to maternal care (oxytocin receptor and estrogen receptor alpha) affected early-life licking received or later-life licking provisioning. However, the effect of these individual SNPs on early-life licking received and later-life licking provisioning was minimal after accounting for multiple comparisons. In addition, we did not genotype the F_0_ parent generation and therefore we were unable to account for the potential effects of the F_0_ mothers’ genotype on early-life licking received. Previous work in mice has demonstrated that the genotype of the offspring can affect the amount of care they receive, termed offspring genetic effects^17,18^, which implies that the genetic profile of an organism can also affect its early-life experience and the organism is not simply a passive recipient of environmental exposures. Additional work is needed to determine biological consequences of these SNPs and it is likely that there are additive effects of different genotypes on maternal care received and provisioning that we were unable to elucidate in this study. The role of genotype in the male lineage may also be an important consideration when studying intergenerational transmission of maternal care, as the fathers provide half their genetic profile to the offspring and play an essential role in genomic imprinting^35^.

We found a gene x environment interaction with average licking received and *Drd2* genotype (rs107017253) on DA levels in the nucleus accumbens, which mediated the indirect association between early-life licking received and later-life licking provisioning. While a mediation analysis typically requires a significant direct association of the independent variable (early-life licking received) on the dependent outcome variable (maternal licking provisioning), we also hypothesized the sequence of causal variables *a priori* based on previous literature on the intergenerational transmission of maternal care^36^. We previously reported similar interactions between average licking received and the same *Drd2* SNP in other DA-related behaviors^21^, but involving the A/A genotype, not A/G. It is possible that the nature of this interaction changes when the brain undergoes significant changes in the dopaminergic system, such as when a female rat is pregnant and preparing for postnatal care of her offspring^37,38^. Identifying other biological mechanisms that underlie these gene x environment interactions, such as linkage disequilibrium and gene expression changes, could help elucidate this discrepancy especially since the female offspring with the A/G genotype are the minority population and therefore may be statistically underpowered in observational studies. Studies that investigate epigenetic mechanisms (e.g., DNA methylation) that would affect the expression of these genes would also be an important component in elucidating mechanisms underlying gene x environment interactions, as both maternal care received and the neural changes that prime mothers to provide care for offspring involve alterations in epigenetic mechanisms and gene expression^4,7,8,39^.

Studies of gene x environment interactions on maternal behavior to date have focused on human cohorts. These studies have identified SNPs in oxytocin-and dopamine-related genes that interact with early-life experiences on different components of mothering behavior^16,40–43^ and that can be mediated by executive function^40^ or depressive symptoms^41^. A strength of our study is application of statistical models typically used in human cohorts to rat populations, where causal biological mechanisms can be more readily examined^44^. More specifically, we can look directly into the rat maternal brain for specific neurotransmitters, neuropeptides, and expression of relevant genes that would not be possible with human populations. This cross-species approach is important to refine hypotheses on factors important for maternal behavior that are evolutionarily conserved.

Overall, this study suggests that offspring genotype for the dopamine receptor 2 gene may be an influential factor when assessing the intergenerational transmission of maternal care. More broadly, offspring genotype in DA-related genes may be a critical factor involved with the development of the dopaminergic system^21,45^. While it has been shown in previous studies that early-life maternal licking received substantially alters the functioning of the dopaminergic system, the underlying biological mechanisms are not as well-understood as other neuroendocrine systems^46^. Elucidating the functional consequences of the *Drd2* SNP analyzed in this study will be important to understand the underlying biological mechanisms in the developmental programming of maternal care and its transmission across generations of female offspring.

## Acknowledgements

This research was supported by operating grants from the Natural Sciences and Engineering Research Council (NSERC) of Canada to ASF and POM.

## Author Contributions

SCL, PP, ASF, and POM designed the study. SCL and PP conducted the maternal care received and maternal care provisioning observations. DGA ran the variant calling analysis from Long-Evans RNA-seq datasets and created the track for the UCSC genome browser. HRM did the brain cryosectioning and liver DNA extractions for genotyping. DC conducted the HPLC. SCL and HRM analyzed the data. ASF and POM and supervised the research. SCL, DGA, HRM, and POM wrote the manuscript. All authors contributed to this manuscript and approved the final version of this manuscript.

## Data Availability

The data that support the findings of this study are available from the corresponding author upon reasonable request.

