## Supplemental Table 1 for "Dopamine Genotype Interacts with Inter-Individual Licking Received on Later-Life Licking Provisioning in Female Rat Offspring"

| **Supplementary Table 1. Publicly available Long-Evans RNA-seq samples used for variant calling.** Data taken from the Gene Expression Omnibus (GEO) and Sequence Read Archive (SRA). Each sample name (SRR) is shown (n = 119), the tissue that the sample was taken from, the sequencing type (RNA-seq or microRNA-seq), and the project that the sample belongs in (n = 5). | | | |
| --- | --- | --- | --- |
| Sample name | Tissue | Sequencing type | Project |
| SRR2155693 | brain | RNA-Seq | PRJNA292611 |
| SRR2155694 | brain | RNA-Seq | PRJNA292611 |
| SRR2155695 | brain | RNA-Seq | PRJNA292611 |
| SRR2155696 | brain | RNA-Seq | PRJNA292611 |
| SRR2155697 | brain | RNA-Seq | PRJNA292611 |
| SRR2155698 | brain | RNA-Seq | PRJNA292611 |
| SRR2155699 | brain | RNA-Seq | PRJNA292611 |
| SRR2155700 | brain | RNA-Seq | PRJNA292611 |
| SRR2155701 | brain | RNA-Seq | PRJNA292611 |
| SRR2155702 | brain | RNA-Seq | PRJNA292611 |
| SRR2155703 | brain | RNA-Seq | PRJNA292611 |
| SRR2155704 | brain | RNA-Seq | PRJNA292611 |
| SRR2155705 | brain | RNA-Seq | PRJNA292611 |
| SRR2155706 | brain | RNA-Seq | PRJNA292611 |
| SRR2927560 | hippocampus | miRNA-Seq | PRJNA301355 |
| SRR2927561 | hippocampus | miRNA-Seq | PRJNA301355 |
| SRR2927562 | hippocampus | miRNA-Seq | PRJNA301355 |
| SRR2927563 | hippocampus | miRNA-Seq | PRJNA301355 |
| SRR2927564 | hippocampus | miRNA-Seq | PRJNA301355 |
| SRR2927565 | hippocampus | miRNA-Seq | PRJNA301355 |
| SRR2927566 | hippocampus | miRNA-Seq | PRJNA301355 |
| SRR2927567 | hippocampus | miRNA-Seq | PRJNA301355 |
| SRR2927568 | hippocampus | miRNA-Seq | PRJNA301355 |
| SRR2927569 | hippocampus | miRNA-Seq | PRJNA301355 |
| SRR2927570 | hippocampus | miRNA-Seq | PRJNA301355 |
| SRR2927571 | hippocampus | miRNA-Seq | PRJNA301355 |
| SRR2927572 | hippocampus | miRNA-Seq | PRJNA301355 |
| SRR2927573 | hippocampus | miRNA-Seq | PRJNA301355 |
| SRR2927574 | hippocampus | miRNA-Seq | PRJNA301355 |
| SRR2927575 | hippocampus | miRNA-Seq | PRJNA301355 |
| SRR2927576 | hippocampus | miRNA-Seq | PRJNA301355 |
| SRR2927577 | hippocampus | miRNA-Seq | PRJNA301355 |
| SRR2927578 | hippocampus | miRNA-Seq | PRJNA301355 |
| SRR2927579 | hippocampus | miRNA-Seq | PRJNA301355 |
| SRR2927580 | hippocampus | miRNA-Seq | PRJNA301355 |
| SRR2927581 | hippocampus | miRNA-Seq | PRJNA301355 |
| SRR2927582 | hippocampus | RNA-Seq | PRJNA301356 |
| SRR2927583 | hippocampus | RNA-Seq | PRJNA301356 |
| SRR2927584 | hippocampus | RNA-Seq | PRJNA301356 |
| SRR2927585 | hippocampus | RNA-Seq | PRJNA301356 |
| SRR2927586 | hippocampus | RNA-Seq | PRJNA301356 |
| SRR2927587 | hippocampus | RNA-Seq | PRJNA301356 |
| SRR2927588 | hippocampus | RNA-Seq | PRJNA301356 |
| SRR2927589 | hippocampus | RNA-Seq | PRJNA301356 |
| SRR2927590 | hippocampus | RNA-Seq | PRJNA301356 |
| SRR2927591 | hippocampus | RNA-Seq | PRJNA301356 |
| SRR2927592 | hippocampus | RNA-Seq | PRJNA301356 |
| SRR2927593 | hippocampus | RNA-Seq | PRJNA301356 |
| SRR2927594 | hippocampus | RNA-Seq | PRJNA301356 |
| SRR4031271 | cortical neurons | RNA-Seq | PRJNA313218 |
| SRR4031272 | cortical neurons | RNA-Seq | PRJNA313218 |
| SRR4031273 | cortical neurons | RNA-Seq | PRJNA313218 |
| SRR4031274 | cortical neurons | RNA-Seq | PRJNA313218 |
| SRR4031275 | cortical neurons | RNA-Seq | PRJNA313218 |
| SRR4031276 | cortical neurons | RNA-Seq | PRJNA313218 |
| SRR4031277 | cortical neurons | RNA-Seq | PRJNA313218 |
| SRR4031278 | cortical neurons | RNA-Seq | PRJNA313218 |
| SRR4031279 | cortical neurons | RNA-Seq | PRJNA313218 |
| SRR4039831 | Lateral entorhinal cortex deep layers | RNA-Seq | PRJNA339395 |
| SRR4039832 | Medial entorhinal cortex layer II | RNA-Seq | PRJNA339395 |
| SRR4039833 | Medial entorhinal cortex deep layers | RNA-Seq | PRJNA339395 |
| SRR4039834 | Lateral entorhinal cortex layer II | RNA-Seq | PRJNA339395 |
| SRR4039835 | Lateral entorhinal cortex deep layers | RNA-Seq | PRJNA339395 |
| SRR4039836 | Medial entorhinal cortex layer II | RNA-Seq | PRJNA339395 |
| SRR4039837 | Medial entorhinal cortex deep layers | RNA-Seq | PRJNA339395 |
| SRR4039838 | Lateral entorhinal cortex layer II | RNA-Seq | PRJNA339395 |
| SRR4039839 | Lateral entorhinal cortex deep layers | RNA-Seq | PRJNA339395 |
| SRR4039840 | Medial entorhinal cortex layer II | RNA-Seq | PRJNA339395 |
| SRR4039841 | Medial entorhinal cortex deep layers | RNA-Seq | PRJNA339395 |
| SRR4039842 | Lateral entorhinal cortex layer II | RNA-Seq | PRJNA339395 |
| SRR4039843 | Lateral entorhinal cortex deep layers | RNA-Seq | PRJNA339395 |
| SRR4039844 | Medial entorhinal cortex layer II | RNA-Seq | PRJNA339395 |
| SRR4039845 | Medial entorhinal cortex deep layers | RNA-Seq | PRJNA339395 |
| SRR4039846 | Lateral entorhinal cortex layer II | RNA-Seq | PRJNA339395 |
| SRR4039847 | Lateral entorhinal cortex deep layers | RNA-Seq | PRJNA339395 |
| SRR4039848 | Medial entorhinal cortex layer II | RNA-Seq | PRJNA339395 |
| SRR4039849 | Medial entorhinal cortex deep layers | RNA-Seq | PRJNA339395 |
| SRR4039850 | Lateral entorhinal cortex layer II | RNA-Seq | PRJNA339395 |
| SRR4039851 | Lateral entorhinal cortex deep layers | RNA-Seq | PRJNA339395 |
| SRR4039852 | Medial entorhinal cortex layer II | RNA-Seq | PRJNA339395 |
| SRR4039853 | Medial entorhinal cortex deep layers | RNA-Seq | PRJNA339395 |
| SRR4039854 | Lateral entorhinal cortex layer II | RNA-Seq | PRJNA339395 |
| SRR4039855 | Lateral entorhinal cortex deep layers | RNA-Seq | PRJNA339395 |
| SRR4039856 | Medial entorhinal cortex layer II | RNA-Seq | PRJNA339395 |
| SRR4039857 | Medial entorhinal cortex deep layers | RNA-Seq | PRJNA339395 |
| SRR4039858 | Lateral entorhinal cortex layer II | RNA-Seq | PRJNA339395 |
| SRR4039859 | Lateral entorhinal cortex deep layers | RNA-Seq | PRJNA339395 |
| SRR4039860 | Medial entorhinal cortex layer II | RNA-Seq | PRJNA339395 |
| SRR4039861 | Medial entorhinal cortex deep layers | RNA-Seq | PRJNA339395 |
| SRR4039862 | Lateral entorhinal cortex layer II | RNA-Seq | PRJNA339395 |
| SRR4039863 | Lateral entorhinal cortex deep layers | RNA-Seq | PRJNA339395 |
| SRR4039864 | Medial entorhinal cortex layer II | RNA-Seq | PRJNA339395 |
| SRR4039865 | Medial entorhinal cortex deep layers | RNA-Seq | PRJNA339395 |
| SRR4039866 | Lateral entorhinal cortex layer II | RNA-Seq | PRJNA339395 |
| SRR4039867 | Lateral entorhinal cortex deep layers | RNA-Seq | PRJNA339395 |
| SRR4039868 | Medial entorhinal cortex layer II | RNA-Seq | PRJNA339395 |
| SRR4039869 | Medial entorhinal cortex deep layers | RNA-Seq | PRJNA339395 |
| SRR4039870 | Lateral entorhinal cortex layer II | RNA-Seq | PRJNA339395 |
| SRR4039871 | Lateral entorhinal cortex deep layers | RNA-Seq | PRJNA339395 |
| SRR4039872 | Medial entorhinal cortex layer II | RNA-Seq | PRJNA339395 |
| SRR4039873 | Medial entorhinal cortex deep layers | RNA-Seq | PRJNA339395 |
| SRR4039874 | Lateral entorhinal cortex layer II | RNA-Seq | PRJNA339395 |
| SRR4039875 | Lateral entorhinal cortex deep layers | RNA-Seq | PRJNA339395 |
| SRR4039876 | Medial entorhinal cortex layer II | RNA-Seq | PRJNA339395 |
| SRR4039877 | Medial entorhinal cortex deep layers | RNA-Seq | PRJNA339395 |
| SRR4039878 | Lateral entorhinal cortex layer II | RNA-Seq | PRJNA339395 |
| SRR4039879 | Lateral entorhinal cortex deep layers | RNA-Seq | PRJNA339395 |
| SRR4039880 | Medial entorhinal cortex layer II | RNA-Seq | PRJNA339395 |
| SRR4039881 | Medial entorhinal cortex deep layers | RNA-Seq | PRJNA339395 |
| SRR4039882 | Lateral entorhinal cortex layer II | RNA-Seq | PRJNA339395 |
| SRR4039883 | Lateral entorhinal cortex deep layers | RNA-Seq | PRJNA339395 |
| SRR4039884 | Medial entorhinal cortex layer II | RNA-Seq | PRJNA339395 |
| SRR4039885 | Medial entorhinal cortex deep layers | RNA-Seq | PRJNA339395 |
| SRR4039886 | Lateral entorhinal cortex layer II | RNA-Seq | PRJNA339395 |
| SRR4039887 | Lateral entorhinal cortex deep layers | RNA-Seq | PRJNA339395 |
| SRR4039888 | Medial entorhinal cortex layer II | RNA-Seq | PRJNA339395 |
| SRR4039889 | Medial entorhinal cortex deep layers | RNA-Seq | PRJNA339395 |
| SRR4039890 | Lateral entorhinal cortex layer II | RNA-Seq | PRJNA339395 |
| SRR4039891 | Lateral entorhinal cortex deep layers | RNA-Seq | PRJNA339395 |
